# *Coxiella burnetii* deubiquitinates host TRAF6 to modulate the macrophage innate immune response

**DOI:** 10.1101/2023.03.15.532774

**Authors:** Arif Sadi, Tatiana M. Clemente, Leonardo Augusto, Rajendra K. Angara, Stacey D. Gilk

**Author notes:** Corresponding author Stacey D. Gilk, University of Nebraska Medical Center 985900 Nebraska Medical Center/DRCII 5031, Omaha, NE 68198-5900. Equal contribution (Arif Sadi, Tatiana M. Clemente). Department of Microbiology and Immunology, Kirksville College of Osteopathic Medicine, A. T. Still University of Health Sciences, Kirksville, MO 63501.

## Abstract

*Coxiella burnetii* is a highly infectious, aerosol-transmitted obligate intracellular bacterium that causes human Q fever and replicates within alveolar macrophages during pulmonary infection. Successful infection requires the *C. burnetii* Type IVB secretion system (T4BSS), which delivers bacterial effector proteins into the host cytoplasm to remodel host processes and suppress the innate immune response. In the lung, IL-17 promotes antibacterial immunity by activating ACT1-TRAF6-dependent pathways that drive inflammatory gene expression, reactive oxygen species production, and neutrophil recruitment. However, how intracellular pathogens subvert IL-17 signaling pathways to promote survival within macrophages remains poorly understood. Here, we show that *C. burnetii* evades IL-17-mediated host defense by targeting TRAF6, a central ubiquitin-dependent regulator of innate immune signaling. IL-17 signaling through ACT1-TRAF6 restricts *C. burnetii* viability in macrophages; however, *C. burnetii* impairs IL-17-induced NF-κB and MAPK activation, particularly JNK phosphorylation. *C. burnetii* T4BSS activity suppresses IL-17-driven reactive oxygen species production and neutrophil recruitment. We identify the *C. burnetii* T4BSS effector EmcB as a TRAF6-targeting deubiquitinase that disrupts IL-17-dependent antimicrobial and chemotactic responses. Together these findings reveal that *C. burnetii* downregulates intracellular innate immune signaling by targeting TRAF6, enabling evasion of the macrophage antimicrobial response and limiting neutrophil-mediated immunity.

**Significance Statement:** *Coxiella burnetii* causes Q fever and survives inside macrophages, immune cells that normally help eliminate inhaled pathogens. This study identifies a mechanism by which *C. burnetii* weakens macrophage defenses. We show that the bacterium disrupts macrophage IL-17 signaling, an immune pathway that promotes inflammatory gene expression, reactive oxygen species production, and neutrophil recruitment. Mechanistically, *C. burnetii* uses the secreted enzyme EmcB to target TRAF6, a central signaling protein required for IL-17-dependent antimicrobial responses. These findings reveal how an intracellular bacterial pathogen disables a key immune signaling hub to promote survival and limit innate immune activation, providing broader insight into how pathogens manipulate host immunity.

## Introduction

*C. burnetii*, the causative agent of human Q Fever, is an obligate intracellular bacterial pathogen that primarily targets alveolar macrophages during natural infection. Intracellularly, *C. burnetii* promotes formation of a phagolysosome-like *<u>C</u>oxiella* <u>C</u>ontaining <u>V</u>acuole (CCV) which supports bacterial replication. Within the CCV, *C. burnetii* uses the specialized Dot/Icm type IVB secretion system (T4BSS) to deliver bacterial proteins into the host cell cytoplasm to manipulate host signaling pathways. Besides maintaining CCV fusogenicity with the endocytic pathway, T4BSS effectors dampen the innate immune response, including blocking apoptosis and inhibiting inflammasome activation^1^. Our previous transcriptome analysis of infected alveolar macrophages identified Interleukin 17 (IL-17) signaling as one of the top pathways manipulated by *C. burnetii* T4BSS effector proteins at early stages of infection^2^. Despite growing evidence that *C. burnetii* suppresses IL-17-dependent host responses^3,4^, how *C. burnetii* and other intracellular pathogens selectively target central innate immune signaling hubs to suppress antimicrobial responses are poorly understood.

IL-17 is a proinflammatory cytokine that plays a key role in protecting the host from infection by both extracellular and intracellular pulmonary pathogens^5^. In the lung, IL-17 is secreted by Th17 cells, γδ T cells and NK T cells, and acts on a variety of target cells due to its ubiquitous receptor^6,7^. In macrophages, IL-17 binding upregulates antimicrobial peptides, chemokine secretion, neutrophil recruitment, and activation of Th1 proinflammatory response, thus leading to pathogen killing^8,9^. IL-17 signals through a dimeric IL-17RA and IL-17RC receptor complex that triggers multiple intracellular signaling pathways through the signaling adaptor Act1 (also known as CIKS - <u>C</u>onnection to <u>IK</u>K and <u>S</u>APK/JNK), which is required for all known IL-17-dependent signaling pathways^10^. Upon ligand binding, Act1 associates with the IL-17 receptor via the conserved SEFIR (Similar Expression of Fibroblast growth factor and IL-17R) domain present in both Act1 and the receptor^11^. Act1 ubiquitinates TRAF6, an adaptor protein and E3 ubiquitin ligase that integrates multiple innate immune signaling pathways to activate the NF-κB and MAPK pathways and induce transcription of inflammatory genes including *IL6, TNFA, CXCL2, CXCL5* and *CCL2*^12^.

In our previous study, we demonstrated that the *C. burnetii* T4BSS downregulates expression of IL-17 host target genes, blocks IL-17-stimulated chemokine secretion, and confers protection from the IL-17 bactericidal effect^2^. While there is evidence that *C. burnetii* T4BSS effector proteins manipulate NF-κB and MAPK signaling pathways^13–15^, upstream pathways such as IL-17 have not been explored. Our *in vitro* data demonstrating that *C. burnetii* inhibits intracellular IL-17 signaling^2^ supports an *in vivo* finding that *C. burnetii*-infected IL-17 receptor knockout mice had a similar bacterial burden as infected wild-type mice^16^. These surprising findings suggest a molecular mechanism whereby one or more *C. burnetii* T4BSS effector proteins target the IL-17 intracellular signaling pathway to downregulate the immune response and promote bacterial survival. Here, we demonstrate that *C. burnetii* targets TRAF6, a central hub integrating innate immune signaling pathways, to subvert IL-17-mediated host defense. By disrupting the IL-17R-ACT1-TRAF6 axis, *C. burnetii* suppresses IL-17-dependent transcriptional responses, macrophage oxidative stress, and neutrophil recruitment. We further identify the T4BSS effector protein EmcB (CBU2013) as a modulator of TRAF6, revealing a strategy by which *C. burnetii* disables a key innate immune signaling hub to suppress antimicrobial defenses and promote intracellular survival.

## Results

### *C. burnetii* suppresses IL-17-ACT1-TRAF6 dependent transcriptional activation

Upon IL-17 receptor engagement, ACT1 recruits TRAF6 and promotes K63-linked TRAF6 polyubiquitination, thereby activating downstream NF-κB and MAPK signaling cascades^17^. Given the central role of this pathway in coordinating pro-inflammatory host-responses and our previous finding that *C. burnetii* targets IL-17 intracellular signaling early during macrophage infection^2^, we hypothesized that *C. burnetii* suppresses IL-17 activation of NF-κB and/or MAPK pathways. Infected alveolar macrophages were stimulated with IL-17 and phosphorylation levels of NF-κB p65, JNK/SAPK, and p38 MAPK were quantified. In mock- and T4BSS-deficient (*ΔdotA*) *C. burnetii* mutant-infected cells, IL-17 stimulation robustly increased phosphorylation of p38 MAPK, and JNK/SAPK, with a modest increase observed for NF-κB p65 (**Figure 1A-C**). In contrast, wild-type *C. burnetii* infection significantly attenuated signaling, reducing p65 and p38 phosphorylation by approximately 40% and 60% (**Figure 1A and B**). Notably, IL-17-induced JNK/SAPK phosphorylation levels were nearly eliminated in WT-infected cells (**Figure 1C**).

**Figure 1.**
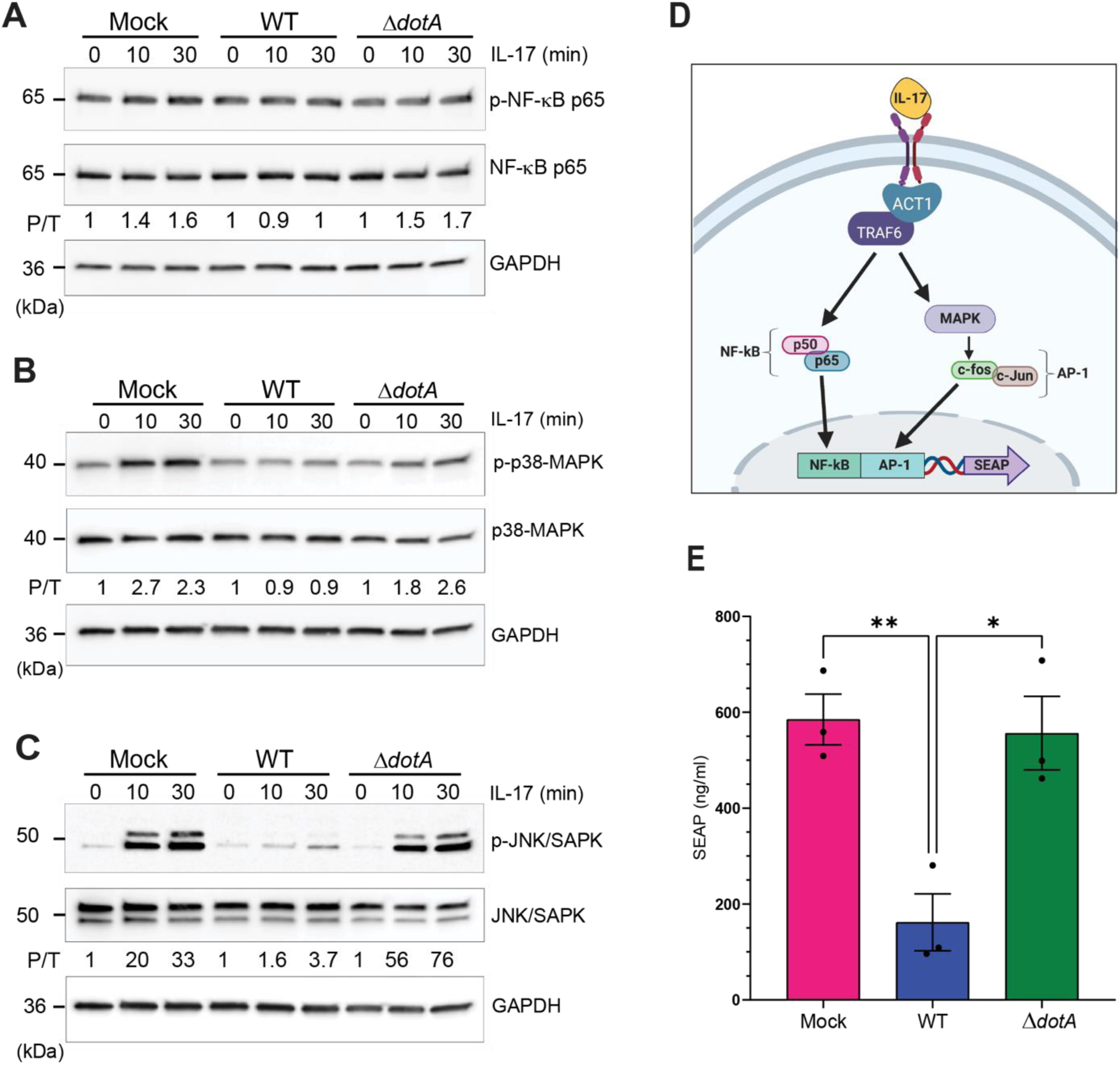
*C. burnetii* suppresses IL-17-ACT1-TRAF6 dependent transcriptional activation. Representative immunoblots of lysates from MH-S cells (alveolar macrophages) either mock-infected or infected with WT or Δ*dotA* mutant *C. burnetii.* Densitometry analysis as indicated by numbers between panels shows decreased phosphorylated levels of **(A)** NF-κB p65 (Ser536), **(B)** p38-MAPK and **(C)** SAPK/ JNK (Thr183/Tyr185) in wildtype-infected cells compared to mock- and Δ*dotA* mutant-infected cells. GAPDH was used as loading control. Following densitometric analysis, the levels of phosphorylated NF-κB, p38 MAPK, and JNK were quantified and normalized to their respective total protein levels and further normalized to values at t=0.Data shown are representative of at least 3 separate experiments. **(D)** HEK-Blue IL-17 SEAP reporter cells stably express the IL-17RA/IL-17RC heterodimer receptor and Act1, along with the secreted embryonic alkaline phosphatase (SEAP) reporter under the control of one NF-κB and five AP-1 (MAPK pathway) binding sites. IL-17 binding to the IL-17 receptor triggers the ACT1-TRAF6 signaling cascade, leading to SEAP transcription. **(E)** HEK-Blue IL-17 SEAP reporter cells were infected for 24h, followed by IL-17 treatment for 24h. SEAP levels were measured using the QUANTI-Blue colorimetric enzyme assay. IL-17 stimulation robustly induced SEAP activity in mock-and *ΔdotA* mutant-infected cells, whereas wild-type *C. burnetii* infection reduced SEAP activity by 72.3%, indicating that *C. burnetii* suppresses IL-17 transcriptional activation. Data shown as means ± SEM from three independent experiments. Statistical significance was determined by one-way ANOVA with Tukey’s multiple comparisons test, *p<0.05, **p<0.01.

We next assessed whether T4BSS effector proteins inhibit IL-17-induced transcription using HEK-Blue™ IL-17 reporter cells (InvivoGen), which stably express IL-17RA/IL-17RC receptor, ACT1, and an NF-κB/AP-1-responsive secreted embryonic alkaline phosphatase (SEAP) reporter^18^. In these reporter cells, IL-17 activates the ACT1-TRAF6 signaling pathway, driving NF-κB and AP-1-dependent SEAP expression (**Figure 1D**). Infected cells were treated with recombinant IL-17, and SEAP activity quantified colorimetrically via p-nitrophenyl phosphate cleavage. Wild-type *C. burnetii* infection reduced IL-17-induced SEAP activity by >70% compared to mock- and T4BSS-defective (*ΔdotA*) mutant-infected cells (**Figure 1E**). Together, these findings establish that the *C. burnetii* T4BSS actively inhibits IL-17-mediated activation of the ACT1-TRAF6 signaling axis, suppressing both kinase activation and downstream transcriptional responses.

### The IL17R-ACT1-TRAF6 pathway is required for IL-17-mediated restriction of *C. burnetii* infection

Our prior work established that IL-17 stimulation enhances macrophage-mediated killing of *C. burnetii*^2^. Given our data that the *C. burnetii* T4BSS suppresses IL-17-dependent activation of the ACT1-TRAF6 pathway, we hypothesized that the IL-17R-ACT1-TRAF6 pathway is required for IL-17-mediated restriction of *C. burnetii* infection. To directly test the requirement for IL-17 receptor and TRAF6-dependent signaling, IL-17RA and TRAF6 macrophage knockouts (*Δil-17ra* and *Δtraf6*) were generated using CRISPR/Cas9 (**Supplemental Figure 1A-B**). Because we were unable to generate an ACT1 knockout in macrophages, siRNA was used to deplete ACT1 protein (**Supplemental Figure 1C**). As expected, IL-17 stimulation of wild-type macrophages reduced *C. burnetii* replication over six days and significantly limited CCV size (**Figure 2A-C**). However, IL-17 treatment failed to restrict bacterial replication or CCV size in *Δil-17ra* and *Δtraf6* macrophages, as well as siRNA-depleted ACT1 macrophages (**Figure 2D and E**). These data indicate that IL-17-mediated antimicrobial activity requires both receptor engagement and downstream ACT1 and TRAF6 signaling.

**Figure 2.**
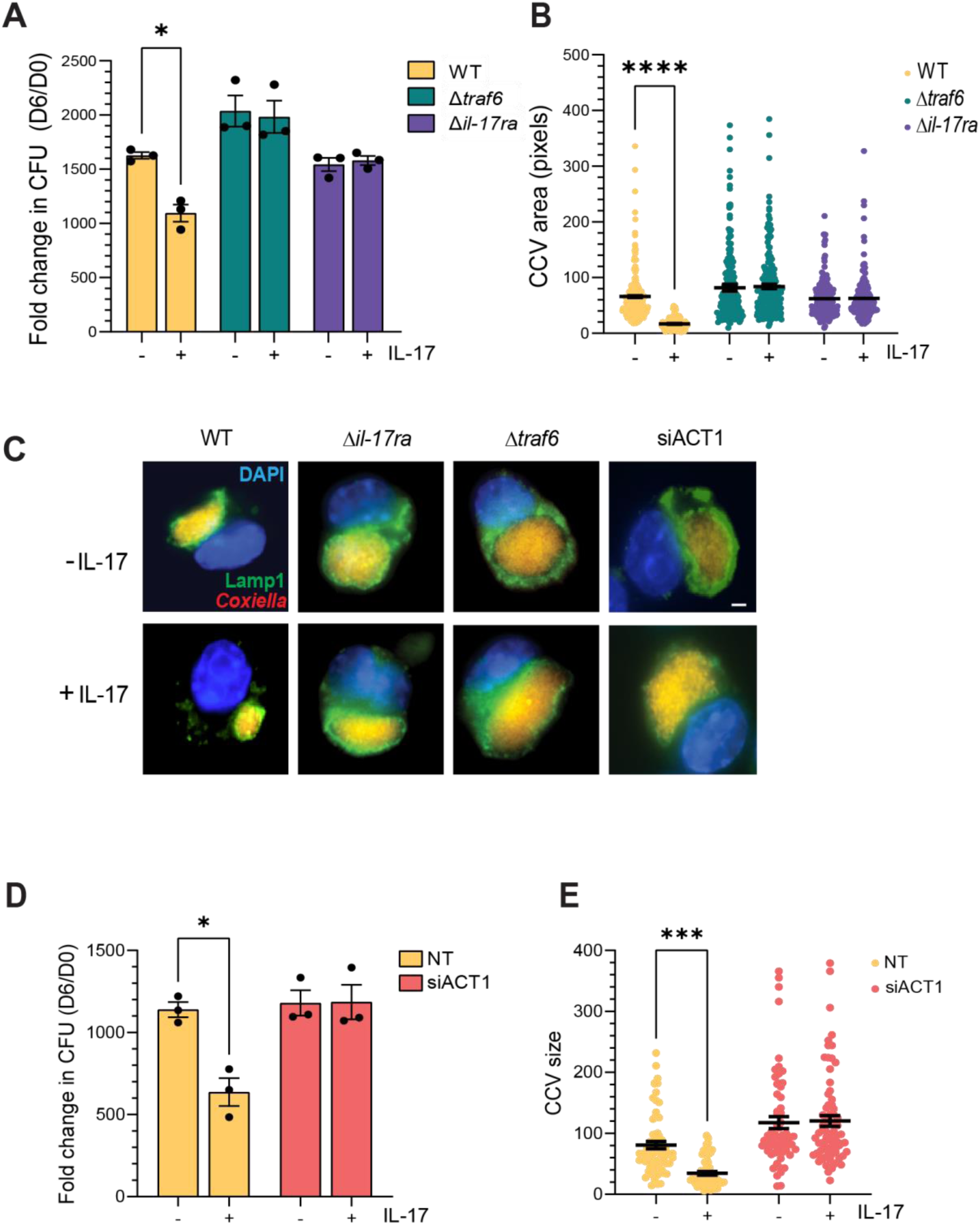
The IL17R-ACT1-TRAF6 pathway is required for IL-17-mediated restriction of *C. burnetii* infection. Stimulation with IL-17 (100 ng/ml) negatively affects *C. burnetii* growth in **(A)** WT MH-S cells or **(D)** control non-targeting siRNA (NT) cells, but not in the **(A)** CRISPR knockouts (*Δil-17ra* and *Δtraf6*) or **(D)** siACT1 MH-S cells, indicating that the bactericidal effect of IL-17 relies on the IL17R-ACT1-TRAF6 pathway. Viable bacteria were quantitated after 6 days using an agarose-based CFU assay, and p values determined by two-way ANOVA, *p<0.05, **p<0.01. Growth was calculated as the fold change over day 0 and shown as the mean ± standard error of the mean (SEM) from three separate growth experiments done in duplicate **(C)** Immunofluorescence staining and **(B and E)** quantitative measurements indicate that CCVs are significantly smaller in WT cells treated with IL-17, but their size is not affected by IL17 in *Δil-17ra*, *Δtraf6*, or siACT1 infected-cells. Representative images of CCVs stained by immunofluorescence at 6 dpi (days post infection; scale bar: 10μm). Blue, DAPI (host cell nuclei); green, LAMP1 (lysosomes and CCV); red, *C. burnetii*. CCV size was measured using ImageJ, with each circle representing an individual CCV. Data are shown as the mean ± SEM of at least 60 cells from three independent experiments. Statistical significance was determined by two-way ANOVA, ****p<0.0001.

Interestingly, *C. burnetii* replication improved in *Δtraf6* macrophages even in the absence of IL-17 stimulation (**Figure 2A**). This IL-17-independent phenotype suggests that TRAF6 may function as a broader innate immune hub restricting *C. burnetii* replication beyond IL-17 signaling alone. Together, these findings establish that the IL-17R-ACT1-TRAF6 axis is required for IL-17-mediated restriction of *C. burnetii* infection and further identify TRAF6 as a key determinant of macrophage antimicrobial defense.

### Targeting TRAF6 inhibits IL-17-induced oxidative stress in macrophages

IL-17 stimulation promotes reactive oxygen species (ROS) production through ACT1-TRAF6-dependent activation of NADPH-oxidases (NOX)^19^. Because elevated ROS restricts *C. burnetii* replication^20^, we hypothesized that *C. burnetii* T4BSS-mediated inhibition of TRAF6 suppresses IL-17-induced oxidative stress in infected macrophages. We first confirmed that IL-17 stimulates NOX activity in alveolar macrophages and observed a significant increase in NOX activity in wild-type macrophages following IL-17 treatment, compared to untreated controls. In contrast, IL-17 failed to increase NOX activity in *Δil-17ra* cells, demonstrating that IL-17-induced ROS required the IL-17 receptor (**Supplemental Figure 2A**).

We next quantified intracellular ROS using CellROX Green in mock-, wild-type- and T4BSS Δ*dotA* mutant-infected cells following IL-17 stimulation. IL-17 treatment significantly increased ROS levels in mock- and Δ*dotA* mutant-infected cells. However, ROS levels did not change in wild-type *C. burnetii*-infected cells (**Figure 3A, B**), indicating that the *C. burnetii* T4BSS suppresses IL-17-induced oxidative stress. To determine whether TRAF6 is required for this response, we measured ROS levels in Δ*traf6* macrophages during infection. In contrast to wild-type macrophages, IL-17 stimulation did not increase ROS levels in *Δtraf6* macrophages, and ROS levels were comparable between mock- and wild-type *C. burnetii*-infected cells (**Figure 3C**). A similar phenotype was observed in Δ*il17ra* macrophages (**Supplemental Figure 2B**), confirming that IL-17-induced oxidative stress requires intact IL-17R-TRAF6 signaling. Together, these findings demonstrate that IL-17-mediated ROS production in macrophages requires TRAF6, and that *C. burnetii* suppresses this antimicrobial oxidative response via its T4BSS. These data further support a model in which bacterial targeting of TRAF6 attenuates IL-17-driven oxidative stress, thereby promoting intracellular survival.

**Figure 3.**
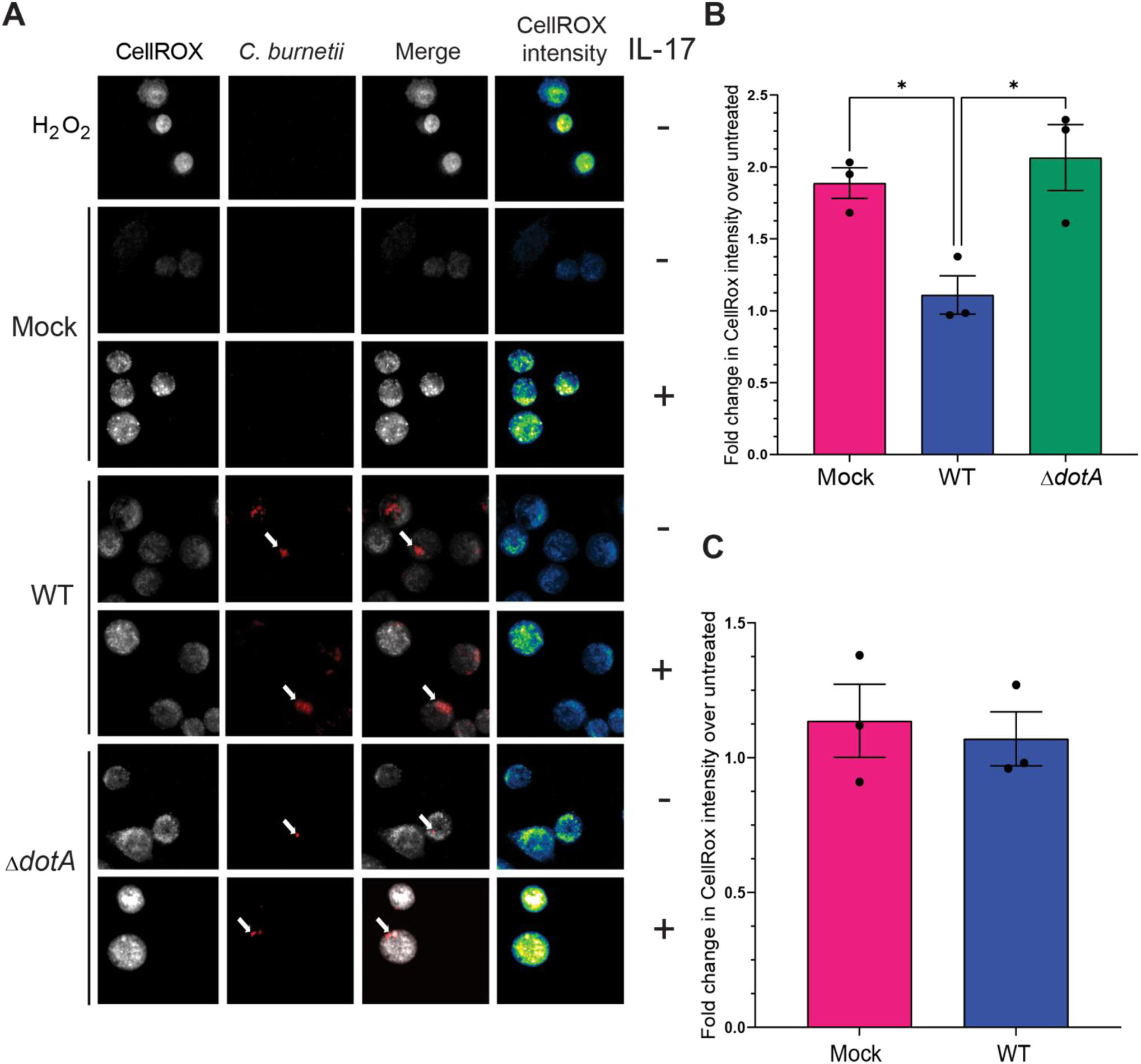
Targeting TRAF6 inhibits IL-17-induced oxidative stress in macrophages. **(A)** MH-S cells (alveolar macrophages) were infected with mCherry-expressing WT or Δ*dotA* mutant bacteria, treated or not with recombinant IL-17 and ROS levels were stained with CellROX Green at 2 dpi. Z-stacks were acquired by live cell spinning disk microscopy. ROS levels are shown as a heat map on the far-right column (CellROX intensity), with yellow showing high levels of ROS and blue showing low levels of ROS (scale bar: 10μm). Arrows indicate bacteria inside cells. **(B)** CellROX Green was used to determine intracellular ROS levels, with fluorescence intensity quantified using ImageJ and normalized to cell area. Stimulation with IL-17 significantly increased the ROS levels in mock- and Δ*dotA* mutant-infected, but not in WT-infected cells, suggesting that *C. burnetii* T4BSS blocks IL-17-induced ROS levels in macrophages. Data is shown as the mean ± SEM of at least 30 cells per condition in each of three independent experiments. Statistical significance was determined by one-way ANOVA with Tukey’s post hoc test, *p<0.05, **p<0.01. **(C)** ROS levels in Δ*traf6* cells. Data are shown as the mean ± SEM of at least 20 cells per condition in each of three independent experiments. Statistical significance was determined Statistical analysis by Two-tailed Unpaired Student’s t-test.

### Targeting TRAF6 inhibits IL-17-dependent chemokine secretion and neutrophil recruitment

IL-17 promotes neutrophil recruitment to the lung by inducing chemokines such as CXCL1, CXCL2, and CXCL5^21^. We hypothesized that *C. burnetii* suppresses TRAF6 signaling to limit IL-17-dependent chemokine secretion, thereby reducing neutrophil recruitment. To first assess chemokine induction, we measured secretion of CXCL5/LPS-induced chemokine (LIX), a potent neutrophil chemoattractant, in *C. burnetii*-infected macrophages following IL-17 stimulation. CXCL5/LIX significantly increased in mock- and Δ*dotA* mutant-infected macrophages after IL-17 treatment (**Figure 4A**). In contrast, IL-17 failed to induce CXCL5/LIX secretion in wild-type *C. burnetii*-infected macrophages, indicating that *C. burnetii* suppresses IL-17-stimulated CXCL5/LIX secretion in a T4BSS-dependent manner.

**Figure 4.**
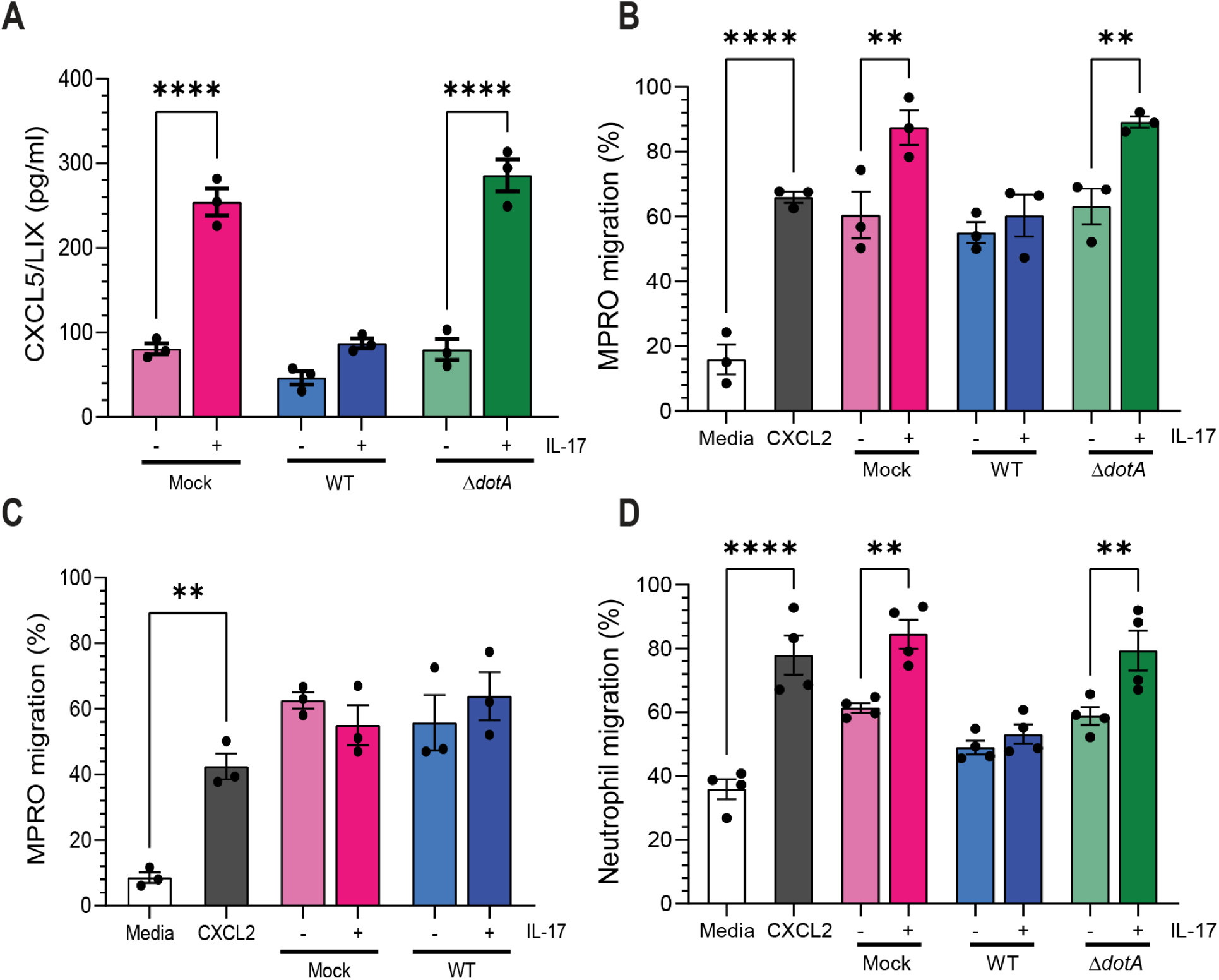
Targeting TRAF6 inhibits IL-17-dependent chemokine secretion and neutrophil recruitment. **(A)** CXCL5/LIX levels were measured in supernatants from mock-, wild-type *C. burnetii*-, or Δ*dotA* mutant-infected macrophages, with or without IL-17 treatment, at 48 hpi. IL-17 stimulated CXCL5/LIX secretion in mock- and Δ*dotA* mutant-infected macrophages, but not in macrophages infected with wild-type bacteria. Statistical significance was determined by one-way ANOVA with Šídák’s multiple comparisons test, ****p<0.0001. **(B)** Mouse promyelocyte-derived neutrophil chemotaxis was measured by transwell migration assay using supernatants from mock-, wild-type *C. burnetii*-, or Δ*dotA* mutant-infected macrophages treated with or without IL- 17. IL-17-treated mock- and ΔdotA mutant-infected macrophage supernatants increased neutrophil chemotaxis, whereas wild-type-infected macrophage supernatants did not. CXCL2 was used as positive control. Statistical significance was determined by one-way ANOVA with Šídák’s multiple comparisons test. **p<0.01 and ****p<0.0001. **(C)** Transwell migration assay of MPRO cells incubated with the supernatant of mock- and WT-infected Δ*traf6* cells treated with IL-17 did not show any difference in neutrophil migration, suggesting *C. burnetii* targets TRAF6 to block IL-17-mediated neutrophil migration. Statistical significance was determined by one-way ANOVA with Šídák’s multiple comparisons test. **p<0.01 **(D)** Primary neutrophils isolated from C57BL/6 bone marrow showed increased chemotaxis toward supernatants from IL-17-treated mock- and ΔdotA mutant-infected macrophages, but not toward supernatants from wild-type-infected macrophages, regardless of IL-17 treatment. Statistical significance was determined by one-way ANOVA with Šídák’s multiple comparisons test., *p<0.05, **p<0.01 and ****p<0.0001. Data is shown as the mean ± SEM from three or four separate experiments.

We next tested whether this defect in chemokine secretion functionally impairs neutrophil recruitment using an *in vitro* chemotaxis assay. CXCL2 served as a positive control for neutrophil migration. Neutrophils differentiated from mouse promyelocyte (MPRO) cells transmigrated robustly toward supernatants from IL-17-treated mock- and *ΔdotA*-infected macrophages, but not toward supernatants from wild-type-infected macrophages, regardless of IL-17 stimulation (**Figure 4B**). These data demonstrate that *C. burnetii* T4BSS-dependent inhibition of IL-17 signaling abrogates macrophage-driven neutrophil chemotaxis. To determine whether TRAF6 is required for this response, we performed parallel chemotaxis assays using supernatants from *Δtraf6* macrophages. In contrast to wild-type macrophages, IL-17 stimulation did not enhance neutrophil migration toward supernatants from either mock- or wild-type-infected *Δtraf6* cells (**Figure 4C**). A similar phenotype was observed in Δ*il17ra* macrophages (**Supplemental Figure 3A**), indicating that IL-17-induced neutrophil migration requires intact IL-17R-TRAF6 signaling. To validate these findings in primary cells, CD11b⁺Ly6G⁺ neutrophils were isolated from murine bone marrow (75.7% purity; **Supplemental Figure 3B**). Consistent with the MPRO results, supernatants from IL-17-treated mock- and *ΔdotA*-infected macrophages induced significant primary neutrophil migration, whereas supernatants from wild-type-infected macrophages did not (**Figure 4D**). Collectively, these data demonstrate that IL-17-dependent neutrophil chemotaxis requires TRAF6 signaling, and that *C. burnetii* suppresses this response via its T4BSS. By targeting TRAF6, the bacterium limits IL-17-induced CXCL5/LIX secretion and downstream neutrophil chemotaxis, thereby dampening host inflammatory cell recruitment.

### EmcB targets TRAF6 to inhibit IL-17-mediated oxidative stress and neutrophil recruitment

Having established the importance of TRAF6 signaling in IL-17-induced oxidative stress and chemotactic responses, we next tested whether *C. burnetii* deubiquitinated TRAF6 during infection in alveolar macrophages (MH-S cells). Endogenous TRAF6 was immunoprecipitated from *C. burnetii*-infected cells and TRAF6-associated ubiquitin levels were assessed by immunoblotting. *C. burnetii* infection reduced TRAF6 ubiquitination by ∼70% compared to uninfected cells **(Figure 5A)**. Because *C. burnetii* suppresses IL-17 signaling in a T4BSS-dependent manner^2^ and TRAF6 ubiquitination is required for IL-17 signal transduction, we next investigated whether a T4BSS effector directly antagonizes TRAF6 ubiquitination. The *C. burnetii* T4BSS effector EmcB (*cbu2013*), recently identified as a ubiquitin-specific cysteine protease^22^, was therefor investigated as a candidate antagonist of TRAF6 ubiquitination. EmcB-mStayGold(mSG) and FLAG-TRAF6 were co-expressed in HEK293T cells, FLAG-TRAF6 was immunoprecipitated, and TRAF6-associated ubiquitin levels were assessed by immunoblotting. EmcB-mSG expression almost completely eliminated TRAF6 ubiquitination compared to mSG alone (**Figure 5B**). In contrast, the catalytic cysteine mutant EmcB C156A, which lacks deubiquitinase activity^22^, did not reduce TRAF6 ubiquitination (**Figure 5B**).

**Figure 5.**
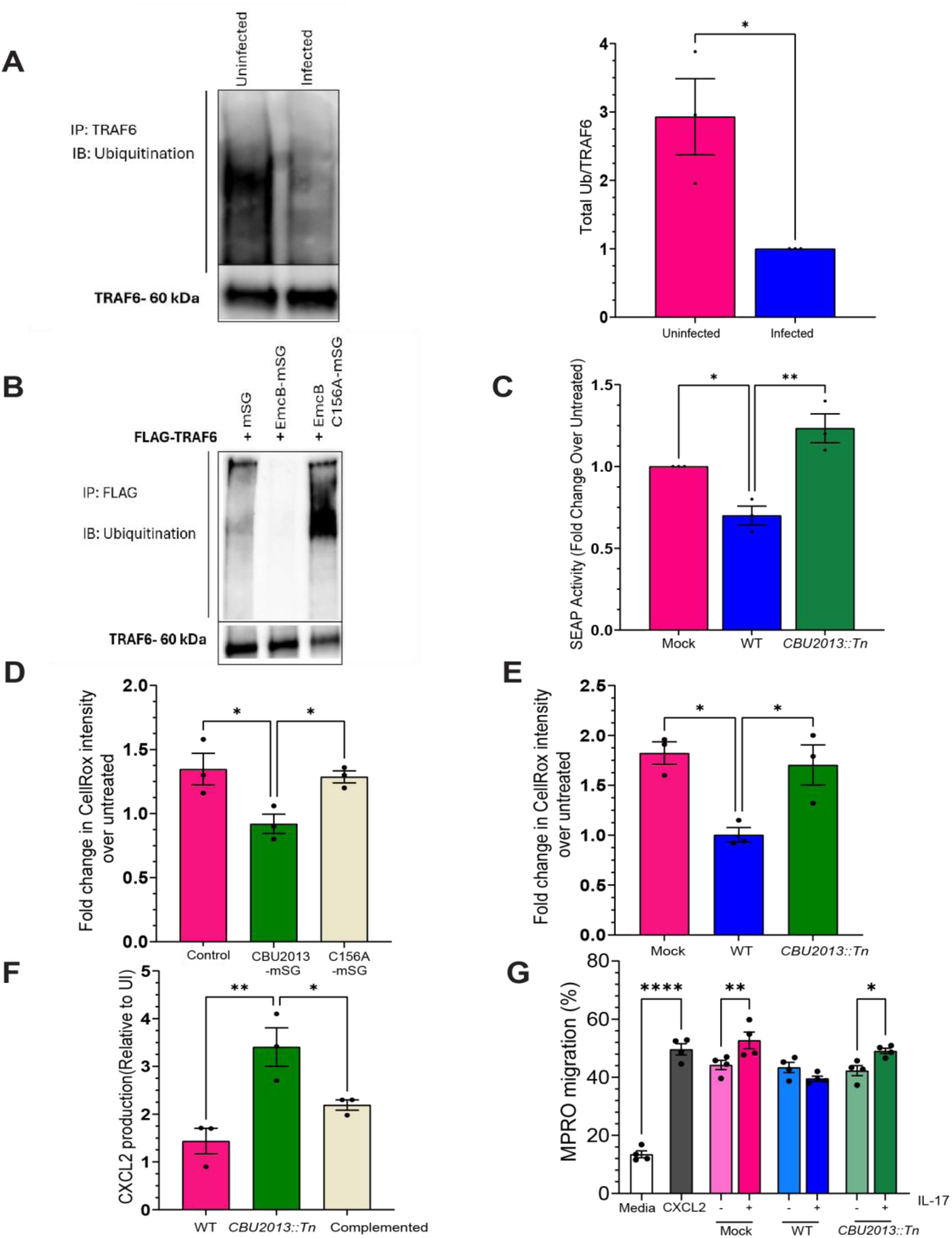
*C. burnetii* T4BSS effector EmcB targets TRAF6 to suppress IL-17 signaling. **(A)** MH-S alveolar macrophages were infected with *C. burnetii* prior to TRAF6 immunoprecipitation and immunoblotting for ubiquitin (left). Total ubiquitination was quantified by densitometry and normalized to TRAF6 levels, with values expressed relative to infected cells (right). Statistical significance was determined using a two-tailed Student’s *t* test, *p<0.05 **(B)** HEK-293T cells were co-transfected with plasmids encoding FLAG-tagged TRAF6 and either the mStayGold vector, EmcB-mStayGold, or C156A-mStayGold. Following immunoprecipitation (IP), samples were subjected to immunoblot (IB) analysis using anti-ubiquitin (top panels) and anti-FLAG (middle panels) antibodies. The presence of mStayGold-linked proteins was identified using an immunofluorescence assay (data not shown). **(C)** SEAP activity was robustly induced upon IL-17 stimulation in both mock-infected and *cbu2013::Tn* (EmcB) mutant-infected HEK-Blue IL-17 reporter cells. In contrast, infection with wild-type *C. burnetii* resulted in a significant reduction in SEAP activity, suggesting that the bacterial effector CBU2013 suppresses IL-17-mediated transcriptional activation. SEAP activity was calculated by comparing IL-17-treated conditions to their corresponding untreated controls, and then normalizing these fold changes to those observed in uninfected cells. Statistical significance was determined by one-way ANOVA with Tukey’s multiple comparisons test, *p<0.05, **p<0.01. **(D)** CellROX Deep Red was used to determine intracellular ROS levels. Cells expressing the catalytically inactive EmcB mutant mSG-C156A exhibited significantly increased ROS following IL-17 stimulation, suggesting that the catalytic cysteine is required for EmcB-mediated suppression of IL-17-induced ROS. Statistical significance was determined by one-way ANOVA with Dunnett’s multiple comparisons test, *p<0.05 **(E)** IL-17 stimulation significantly increased ROS levels in mock- and *cbu2013::Tn* mutant-infected cells, whereas wild-type *C. burnetii* infection prevented this response, suggesting that EmcB suppresses IL-17-induced ROS production in macrophages. Data is shown as the mean ± SEM of at least 20 cells per condition in each of three independent experiments. Statistical significance was determined by one-way ANOVA with Tukey’s post hoc test, *p<0.05 **(F)** CXCL2 levels were quantified by ELISA in supernatants of macrophages infected wild-type *C. burnetii*, the *cbu2013::Tn* mutant, or the the complemented mutant at 48 hours post infection. Loss of EmcB increased CXCL2 secretion, whereas complementation restored suppression of this IL-17 target chemokine. Statistical significance was determined by one-way ANOVA with Dunnett’s multiple comparisons test, *p<0.05, **p<0.01 **(G)** MPRO-derived neutrophil chemotaxis was measured using supernatants from mock-, wild-type *C. burnetii*-, or *cbu2013::Tn* mutant-infected macrophages treated with or without IL-17. Supernatents from IL-17-treated mock- and *cbu2013::Tn* mutant-infected macrophage increased neutrophil chemotaxis, whereas wild-type-infected macrophage supernatants did not. CXCL2 was used as a positive control. CXCL2 was used as positive control. Statistical significance was determined by one-way ANOVA with Šídák’s multiple test, *p<0.05, **p<0.01 and ****p<0.0001. Data is shown as the mean ± SEM from three or four separate experiments.

Because K63-linked ubiquitination of TRAF6 is required for IL-17-dependent activation of NF-κB and MAPK signaling, we next tested whether EmcB suppresses IL-17-dependent transcription. HEK-Blue™ IL-17 reporter cells were infected with wild-type or *cbu2013::Tn* mutant *C. burnetii*^23^ and IL-17-induced SEAP activity measured. Cells infected with a *C. burnetii cbu2013::Tn* mutant exhibited significantly higher IL-17-induced SEAP activity compared to cells infected with wild-type *C. burnetii* (**Figure 5C**). These data indicate that EmcB is required to suppress IL-17-mediated transcription activation during *C. burnetii* infection. Collectively, these data identify TRAF6 as a host target of the *C. burnetii* T4BSS effector EmcB and provide a mechanistic basis for bacterial inhibition of IL-17 signaling through deubiquitination of a central inflammatory signaling hub.

Having established that EmcB deubiquitinates TRAF6 and suppresses IL-17-dependent transcription, we next asked whether this was sufficient to inhibit downstream antimicrobial effector responses. We first quantified IL-17-induced ROS generation in cells ectopically expressing wild-type EmcB or a catalytically inactive EmcB mutant (C156A). Following IL-17 stimulation, untransfected cells and cells expressing EmcB C156A exhibited significantly elevated ROS levels **(Figure 5D)**. In contrast, cells expressing wild-type EmcB showed markedly reduced ROS production, indicating that the deubiquitinase activity of EmcB is required to suppress IL-17-induced oxidative stress. We next determined whether EmcB mediates ROS suppression during infection. Intracellular ROS levels were quantified using CellROX Deep Red in mock-, wild type-, and *cbu2013::Tn*-infected macrophages following IL-17 stimulation. IL-17 treatment significantly increased ROS production in mock- and *cbu2013::Tn*-infected cells, whereas wild type-infected macrophages failed to mount an oxidative response **(Figure 5E)**. Thus, EmcB is required for *C. burnetii* suppression of IL-17-induced ROS during infection.

Because TRAF6 signaling also promotes IL-17-dependent chemokine production, we next examined whether EmcB suppresses chemokine secretion and the resulting neutrophil recruitment. CXCL2 secretion was measured by ELISA in macrophages infected with wild-type *C. burnetii*, the *cbu2013::Tn* mutant, or the complemented *cbu2013::Tn* strain. Macrophages infected with the *cbu2013::Tn* mutant secreted significantly more CXCL2 than those infected with wild-type bacteria, whereas genetic complementation restored suppression (**Figure 5F**). These data demonstrate that EmcB is required for *C. burnetii*-mediated suppression of chemokine production. Finally, we assessed the functional consequence of EmcB-mediated chemokine suppression using an *in vitro* neutrophil chemotaxis assay. Recombinant CXCL2 robustly induced neutrophil migration, confirming assay sensitivity. Supernatants from mock- and *cbu2013::Tn*-infected macrophages stimulated with IL-17 significantly increased neutrophil migration, whereas supernatants from wild-type-infected macrophages failed to promote migration **(Figure 5G)**. Collectively, these findings establish EmcB as a *C. burnetii* T4BSS-secreted deubiquitinase that targets TRAF6 to suppress IL-17-driven antimicrobial signaling, thereby suppressing multiple IL-17-dependent innate immune effector responses, including macrophage ROS production, chemokine secretion, and neutrophil recruitment.

## Discussion

The innate immune response relies on proinflammatory cytokines to trigger and regulate an effective response to invading pathogens. The proinflammatory cytokine IL-17 promotes protective innate immunity by inducing antimicrobial peptides, stimulating ROS production, recruiting neutrophils, and promoting IFN-γ responses^24^. Our previous studies revealed that *C. burnetii* downregulates intracellular IL-17 signaling in macrophages in a T4BSS-dependent manner, implicating one or more T4BSS secreted effector proteins^2^. Here, we demonstrate that IL-17-mediated restriction of *C. burnetii* requires IL-17R, ACT1, and TRAF6, and that *C. burnetii* disrupts this pathway by promoting TRAF6 deubiquitination, thereby attenuating NF-κB and MAPK signaling. We further show that EmcB, a *C. burnetii* T4BSS effector and deubiquitinase, targets TRAF6 to inhibit IL-17-mediated oxidative stress and IL-17-induced neutrophil chemotaxis. Together, these findings identify TRAF6 as a pathogen-targeted signaling hub in the IL-17 antimicrobial response and define EmcB as a bacterial effector that disrupts TRAF6-dependent oxidative and chemotactic responses.

Alveolar macrophages play a key role in immune surveillance of the airway and phagocytosis of inhaled bacteria, limiting pathogen replication while preserving pulmonary function^25^. However, intracellular bacterial pathogens, including *Mycobacterium tuberculosis* and *Yersinia pestis*, have evolved mechanisms to subvert macrophage antimicrobial responses, including inflammatory signaling and oxidative stress pathways^26,27^. One common mechanism to evade the early host immune response is by modulating key inflammatory signaling pathways. We found that *C. burnetii* blocks IL-17-induced activation of NF-κB and MAPK pathways, dampening the immune response. Although *C. burnetii* has previously been shown to modulate NF-κB and MAPK in other contexts^13–15^, this study identifies the upstream IL-17R-ACT1-TRAF6 axis as a specific pathway targeted during infection.

IL-17 signaling depends on TRAF6 E3 ubiquitin ligase activity and K63-linked TRAF6 ubiquitination, which creates non-degradative ubiquitin scaffolds that activate downstream signaling complexes, including TAK1, NF-κB, and MAPK ^28,29^. Our finding that EmcB deubiquitinates TRAF6 provides a mechanistic explanation for how *C. burnetii* attenuates the IL-17 signaling pathway. Given prior evidence that EmcB preferentially cleaves K63-linked ubiquitin chains^22^, it is possible that EmcB removes K63-linked polyubiquitin chains from TRAF6. Consistent with this model, EmcB-mediated deubiquitination of TRAF6 is strictly dependent on catalytic activity, as this effect is abolished by mutation of the catalytic cysteine. Collectively, these findings support a model in which EmcB targets TRAF6-dependent signaling, likely by removing ubiquitin scaffolds required for downstream NF-κB and MAPK activation.

Pathogen-mediated disruption of TRAF6 ubiquitin signaling is not unique to *C. burnetii*. Host deubiquitinating enzymes (DUBs), such as A20, CYLD and USP family members, modulate NF-κB signaling by removing ubiquitin chains from TRAF6^30^. Bacterial pathogens have evolved analogous strategies to inhibit inflammatory signaling. The *Yersinia* Type III effector protein YopJ inhibits TRAF6 and TRAF2 ubiquitination to block p38 MAPK, JNK, and NF-κB activation^31^, while the *Shigella flexneri* effector OspI modifies UBC13, thereby inhibiting TRAF6 activation and inflammatory signaling^32^. EmcB appears to act through a distinct biochemical mechanism with direct deubiquitination of TRAF6, but converges on the same host vulnerability: the requirement for TRAF6 ubiquitin scaffolds in innate immune signaling. Prior work identified EmcB as a deubiquitinase that targets RIG-I and suggested that the DUB has limited substrate specificity based on minimal changes in global K63-linked ubiquitination^22^. However, deubiquitinases can act on multiple substrates since they recognize ubiquitin itself in addition to the modified protein substrate^33^. Recent structural analyses further suggest that the EmcB deubiquitinase domain may allow recognition of multiple host substrates^34^.

IL-17 activates NADPH oxidases in an ACT1- and TRAF6-dependent manner^19^. Our data indicate that *C. burnetii* targets the IL-17R-ACT1-TRAF6 pathway to block ROS production, likely as a strategy to promote bacterial survival. This interference highlights TRAF6 as a critical regulatory node that pathogens may exploit to dampen host oxidative defenses. Such a strategy is not without precedent, as the *Salmonella* Typhimurium effector protein SopB inhibits mitochondrial recruitment of TRAF6, effectively reducing mitochondrial ROS generation^35^. In this context, TRAF6 emerges as a convergent target for distinct intracellular pathogens seeking to evade ROS-mediated killing. *C. burnetii* may also employ additional mechanisms to counteract ROS production. For example, RpoS, a stationary-phase sigma factor required for bacterial survival during environmental stress, plays an important role in ROS resistance^36^. In addition, a recent study characterized the *C. burnetii* effector protein SdrA as a short-chain dehydrogenase, which is also essential for bacterial resistance to oxidative stress and intracellular replication^37^.

In addition to modulating ROS, TRAF6 regulates IL-1, TLR4, IL-17 signaling pathways that contribute to neutrophil recruitment^38–40^. Consistent with this role, cells infected with an EmcB-deficient strain had increased CXCL2 secretion and IL-17-induced neutrophil chemotaxis, suggesting that EmcB-dependent suppression of TRAF6 signaling limits production of neutrophil-recruiting factors. This finding may be relevant *in vivo*, as neutrophil recruitment to the lungs is delayed until approximately seven days post-infection following aerosol *C. burnetii* infection, despite evidence that neutrophils contribute to protective immunity^16,41^. Thus, EmcB-mediated suppression of IL-17-dependent chemokine production provides a plausible mechanism by which *C. burnetii* may delay neutrophil recruitment during early pulmonary infection.

EmcB-mediated TRAF6 deubiquitination may have broader consequences for *C. burnetii* infection. TRAF6 is involved in autophagy, inflammasome activation, and the adaptive immune response, and TRAF6-mediated ubiquitination of BECN1/Beclin 1 and Rab7 has been implicated in autophagy induction and autophagosome-lysosome fusion^42–44^. Because EmcB has also been linked to autophagy^45^, TRAF6 deubiquitination could contribute to additional immune defects during *C. burnetii* infection. These possibilities remain to be tested but suggest that EmcB may broadly remodel TRAF6-dependent host defense pathways.

Although our data support a model in which EmcB disrupts TRAF6-dependent IL-17 signaling, defining the specific ubiquitin linkages removed from TRAF6 during infection will be important. In addition, EmcB has been implicated in targeting other ubiquitinated host proteins, including RIG-I, raising the possibility that its effects on macrophage activation reflect coordinated regulation of multiple immune substrates. Future studies using linkage-specific ubiquitin approaches, substrate-trapping EmcB mutants, and infection models expressing catalytically inactive EmcB will be important to determine how TRAF6 deubiquitination contributes to pathogenesis *in vivo*.

Together, our findings identify TRAF6-dependent ubiquitin signaling as a key vulnerability in the IL-17 antimicrobial response and reveal EmcB as a bacterial effector that exploits this node to suppress macrophage ROS production and neutrophil recruitment. These results expand the known functions of *C. burnetii* T4BSS effectors and support a model in which targeted disruption of TRAF6 allows *C. burnetii* to blunt early innate immune responses in the lung.

## Materials and Methods

### Bacteria

*C. burnetii* Nine Mile phase II (NMII clone 4, RSA 439) mCherry-wild type (WT), mCherry- Δ*dotA* mutant^46^ and GFP-*cbu2013::Tn*^23^ were grown for 8 days in ACCM-2, washed twice with PBS and stored as previously described^47^. The GFP-*cbu2013::Tn* strain was complemented in single copy using the *cbu2013* gene under the p1169 promoter (pJB-2HA-CBU2013). For transformation, an 8 day ACCM-D culture of *cbu2013::Tn* was pelleted (16,000×g, 15 min, 4°C), washed, resuspended in ice-cold 10% glycerol, and electroporated with pJB-2HA-CBU2013 (1.8kV, 500Ω, 25µF). Transformants were selected with kanamycin and chloramphenicol and verified by immunoblotting using an anti-HA antibody (Thermo Scientific cat # 26183).

### Mammalian cells and infection

Murine alveolar (MH-S) macrophages (CRL-2019; ATCC) and HeLa cells (CCL-2; ATCC) were maintained in RPMI 1640 (Corning) containing 10% FBS (Atlanta Biologicals). HEK293T cells (ATCC CRL-3216) and IL-17 promoter reporter cells (HEK-Blue™ IL-17 cells; InvivoGen) were cultured in DMEM with high glucose (Corning) with 10% FBS. The multiplicity of infection (MOI) was optimized for each bacterial stock and culture vessel for a final infection level of approximately 1 internalized bacterium per cell.

MPRO clone 2.1 cells (CRL-11422; ATCC) were maintained in Iscove’s Modified Dulbecco’s Medium (IMDM; ATCC) supplemented with 20% equine serum (HyClone, Cytiva) and 10 ng/ml of GM-CSF (BioLegend). MPRO cells were differentiated into neutrophils using 10μM all-trans-Retinoic acid (ATRA; Thermo Scientific Chemicals), for 3 days^48^.

To obtain primary neutrophils, mouse bone marrow cells were isolated from tibia and femurs removed from male C57BL/6 mice, as previously described in^49^. Neutrophils were negatively selected using the MojoSort™ Mouse Neutrophil Isolation Kit Protocol (BioLegend) and immediately used for the migration assays. Neutrophil purity was confirmed using flow cytometry for CD11b^+^ and Ly6G^hi^ (BioLegend). Briefly, 2 × 10⁶ cells were pelleted by centrifugation at 150 × g for 5 min and resuspended in 465 µL of PBS supplemented with 0.1% serum. Five aliquots containing approximately 4 × 10⁵ cells (92.5 µL per tube) were prepared. Each aliquot was incubated with 2.5 µL of anti-CD16/32 antibody (FcγRII/III) (BioLegend 101302) for 5 min at room temperature to block Fc receptors. Fluorophore-conjugated antibodies were added, and samples were incubated for 30 min at 4°C. Following incubation, 1 mL of PBS/serum was added to each tube, and cells were pelleted by centrifugation at 7,000 rpm for 1 min. Supernatants were aspirated, and cell pellets were resuspended in 300 µL of PBS/serum before transfer to flow cytometry tubes.

### IL-17 SEAP reporter assay

HEK-Blue™ IL-17 cells were plated in a 6-well plate (2×10^5^ cells per well), and infected with *C. burnetii* in 0.5 ml growth medium for 2 h at 37°C in 5% CO_2_. Infected cells were washed with PBS and incubated in 2 ml growth medium. At 24 hours-post infection (hpi), cells were treated with 100 ng/ml human IL-17 recombinant protein (R&D Systems). At 48 hpi, the supernatant was collected and 20 μl added to white 96-well plate with transparent bottom. The secreted embryonic alkaline phosphatase (SEAP) was measured in a microplate reader (OD Ex/Em=620/655 nm) using Quanti-Blue Solution (InvivoGen) following the manufacturer’s instructions. A standard curve was generated by plotting absorbance (colorimetric) as a function of SEAP concentration and fitted using linear regression within the linear range of the assay. The standard curve was used to determine SEAP concentration of the samples.

### Phospho-protein immunoblotting

MH-S cells were plated in a 6-well plate (2×10^5^ cells per well) and allowed to adhere overnight. Cells were either mock-infected or infected with *C. burnetii* in 0.5 ml RPMI 10% FBS for 2 h. Infected cells were washed with PBS and incubated in 2 ml of fresh growth medium. At 24 hpi, cells were stimulated with 100 ng/ml mouse IL-17 recombinant protein (R&D Systems) or vehicle control for 10 or 30 min. Cells were lysed with RIPA buffer (Cell Signaling Technologies) containing phosphatase and protease inhibitors (Sigma-Aldrich) and protein lysates resolved by 4-20% SDS-PAGE and transferred to nitrocellulose membrane (BioRad). The membrane was blocked in 5% milk in TBS-T (TBS containing 0.05% tween-20), for 1 h, and probed separately using the following primary rabbit antibodies (1:1000, Cell Signaling Technologies): anti-NF-κB p65 (Cell Signaling 8242), anti-phospho NF-κB p65 (Ser536)(Cell Signaling 3033), anti-SAPK/JNK (Cell Signaling 9252), anti-phospho SAPK/JNK (Thr183/Tyr185) (Cell Signaling 4688), anti-p38 MAPK (Cell Signaling 9212) or anti-phospho p38 MAPK (Cell Signaling 9211) in 5% milk in TBS-T. GAPDH was probed as a loading control (1:1000; Thermo Fisher Scientific MA5-15738). After washing, the membrane was incubated for 1 h with the secondary-antibody horseradish peroxidase (HRP)-conjugated anti-rabbit or anti-mouse (1:1000; Thermo Fisher Scientific) in 5% milk in TBS-T, washed and developed using enhanced chemiluminescence (ECL) reagent (SuperSignal West Pico PLUS; Thermo Fisher Scientific). Densitometry quantitation was performed in ImageJ (Fiji) software using the Easy Band Quantification plugin^50^; the levels of phosphorylated NF-κB, p38 MAPK, and JNK were normalized to their respective total protein levels, and the resulting phosphorylation-to-total (P/T) ratios compared across conditions and normalized to values at t=0.

### CRIPSR knockout macrophages

Disruption of the *il17ra* and *traf6* genes in MH-S cells was carried out using CRISPR/Cas9^51^. sgRNA (IL-17RA: GCTCTGCACCCTCGAGGTAC; TRAF6: ATTTGGGCACTTTACCGTCA) were complexed with EnGen Spy Cas9 NLS protein (New England BioLabs) and transfected into MH-S cells (Lonza, P3 Primary Cell 4D-Nucleofector X kit). After 48 h, the cells were cloned using a 96-well plate. *il17ra* and *traf6* clones were validated by RT-qPCR using Syber green. *traf6* forward primer 5’-CCTTTGGCAAATGTCATCTGTG-3’ and reverse primer 5’-CTCTGCATCTTTTCATGGCAAC3-’; *il17ra*forward primer 5’ - CCCAGTAATCTCAAATACCACAGTTC3-’ reverse primer 5’-GATGAGTGTGATGAGGCCATA-3’.

### Protein depletion with siRNA

MH-S cells (3×10^5^ cells/well in a 6-well plate) were reverse transfected with 50 nM small-interfering RNA (siRNA) SMARTpool specific for mouse ACT1 or non-targeting (NT) control using DharmaFECT 4 Transfection Reagent (Horizon Discovery). After 48 h, cells were infected for 2 h. Following washing with PBS, cells were harvested by trypsinization and subjected to a second round of siRNA reverse transfection in 24-well plate (3.5×10^4^ cells/well). At 2- and 5-days post-infection (dpi), cells were harvested, lysed with 1x RIPA Buffer containing phosphatase and protease inhibitors, and analyzed by immunoblotting to confirm ACT1 knockdown using mouse ACT1 (1:1000; Santa Cruz sc-398161) and mouse GAPDH (1:1000; Thermo Fisher Scientific MA5-15738) antibodies in 5% milk in TBS-T, with GAPDH as the loading control. After washing, the blot was incubated with HRP-conjugated anti-mouse secondary antibody in 5% milk in TBS-T and developed using enhanced chemiluminescence (ECL) reagent. Densitometry analysis was performed using the ImageJ Easy Band Quantification plugin, normalizing to GAPDH to determine knockdown efficiency^50^.

### *C. burnetii* intracellular growth assay

Wildtype, *Δil-17ra* and *Δtraf6* MH-S cells were plated in a 6-well plate (2×10^5^ cells per well) and allowed to adhere overnight; siACT1 and siNT cells were transfected 2 days before infection as described above. All cells were infected with wildtype *C. burnetii* in 0.5 ml RPMI for 2 h, washed with PBS, and scraped into 2 ml of fresh growth medium. Infected cells were replated in a 24-well plate (3×10^4^ cells per well), with siNT and siACT1 cells subjected to a second round of siRNA reverse transfection, replating onto a 24-well plate (4×10^4^ cells/well). Cells were stimulated with 100 ng/ml recombinant mouse IL-17 and the media was changed daily to ensure constant IL-17 concentration. To determine the number of internalized bacteria at days 0 and 6, infected cells were lysed in sterile water for 5 min, diluted in ACCM-2 and spotted 0.25% ACCM-2 agarose plates^52^. The plates were incubated for 7 to 9 days and colonies counted to measure bacterial viability, normalizing to day 0 to determine fold change in growth. Each of the three experiments was performed in biological duplicate, and the bacteria were spotted in triplicate.

### CCV size

Wildtype, *Δil-17ra* and *Δtraf6* MH-S cells were plated in a 6-well plate (2×10^5^ cells per well) and allowed to adhere overnight. All cells were infected with mCherry-expressing wild type *C. burnetii* in 0.5 ml RPMI for 2 h, washed with PBS, and scraped into 2 ml of fresh growth medium. Infected cells were replated onto coverslips in a 24-well plate (3×10^4^ cells per well). Cells were stimulated with 100 ng/ml recombinant mouse IL-17 protein, and the media was changed daily with fresh IL- 17. At 6 dpi, cells were fixed with 2.5% paraformaldehyde (PFA) for 15 min, washed in PBS, and blocked/permeabilized in 1% BSA and 0.1% saponin in PBS for 20 min. Coverslips were stained with rat anti-mouse LAMP1 (1:1,000; BD Biosciences 553792) along with guinea pig anti-*C. burnetii* (1:2500) for 1 h followed by Alexa Fluor secondary antibodies (1:1,000; Invitrogen) for 1 h. Following washing with PBS, coverslips were mounted with ProLong Gold with DAPI and visualized on a Nikon eclipse Ti2 microscope using a 60x oil immersion objective. Images were captured and processed identically, and the CCV area was measured using ImageJ (Fiji) software. At least 30 CCVs were measured per condition for each experiment.

### NADH Oxidase activity

Wildtype and *Δil-17ra* MH-S cells were plated in a 6-well plate (2×10^5^ cells per well). The next day, IL-17 (100 ng/ml) was added and NOX activity quantitated 24 h later using the NADH Oxidase Activity Assay Kit (Abcam) following the manufacturer’s protocol. Briefly, this fluorometric assay links oxidation-reduction reactions of a colorless probe, yielding a brightly colored product that fluoresces at excitation/emission wavelengths of 535/587 nm. The fluorescence generated is directly proportional to the NOX activity in samples. Standard curve generation and sample processing were performed as follows: the 0 standard reading was subtracted from all standard values prior to plotting the lactate standard curve; blank readings were subtracted from all sample readings and corrected sample values were interpolated from the lactate standard curve to determine product formation. NOX activity was calculated using the equation: NOX activity = B / (ΔT × P) where B represents the lactate amount derived from the standard curve (pmol), ΔT corresponds to the reaction time (T₂ − T₁, in minutes), and P denotes the total protein content of the sample (mg). Results are expressed as pmol/min/mg (μU/mg).

### ROS measurements

For experiments in MH-S cells, wildtype, *Δil-17ra* and *Δtraf*6 cells were plated in a 6-well plate (2×10^5^ cells per well) and allowed to adhere overnight. Cells were infected with mCherry-expressing WT or *ΔdotA* mutant *C. burnetii* in 0.5 ml RPMI 10% FBS, for 2 h. Infected cells were washed with PBS, trypsinized, resuspended to 3 x 10^5^ cells/ml and plated onto ibidi-treated channel μ-slide VI0.4 (9×10^3^ cells per channel; Ibidi). After 24 h, the infected cells were treated with 100 ng/ml recombinant mouse IL-17 protein or vehicle control. At 2 dpi, the cells were treated with 10 μM hydrogen peroxide (Thermo Fisher Scientific) for 2 h, and then incubated with 5 μM CellROX Green (Thermo Fisher Scientific) for 30 min. After washing with PBS, cells imaged in growth medium using z-stacks of 0.3-μm step with a Nikon spinning disk confocal microscope (60x oil immersion objective). Images were captured and processed identically, and fluorescence intensity measured using ImageJ (Fiji) software, normalizing to the corresponding untreated control. CellROX DeepRed (Thermo Fisher Scientific) was used when cells were infected with GFP-expressing WT or *cbu2013::Tn* mutant *C. burnetii*.

HeLa cells were reverse-transfected with mStayGold, EmcB-mStayGold and EmcB-mStayGold (C156A) using FuGENE6 (Promega) according to the manufacturer’s protocol. 48 h later, cells were trypsinized, resuspended to 3 x 10^5^ cells/ml, and plated onto ibidi-treated channel μ-slide VI0.4 (9 × 10^3^ cells per channel; Ibidi). The cells were then treated or not with 100 ng/ml human IL-17 recombinant protein for 2h. and then incubated with 5 μM CellROX Deep Red (Thermo Fisher Scientific) for 30 min. After washing with PBS, cells were incubated in growth medium and imaged live using z-stacks of 0.3-μm steps with a Nikon spinning disk confocal microscope (60x oil immersion objective). Images were captured and processed identically; fluorescence intensity was measured using ImageJ (Fiji) software. ROS measurements (fluorescence intensity) were normalized to the corresponding untreated control.

### ELISA

CXCL2-MIP-2 protein levels in cell-free supernatants were measured by ELISA (R&D Systems) according to the manufacturer’s instructions. In brief, MH-S cells (5 × 10^4^ cells per well of a 24-well plate) were plated and allowed to adhere overnight. The cells were then mock infected or infected *C. burnetii* in 0.25 ml growth medium for 2 h at 37°C in 5% CO_2_, washed with PBS, and incubated in 0.5 ml growth medium. At 24 hpi, cells were stimulated or not with 100 ng/ml mouse IL-17 recombinant protein. The cell supernatant was collected at 48 hpi, centrifuged at 20,000 × *g* for 10 min, and analyzed by ELISA.

### Multiplex cytokine profiling assay

MH-S cells were plated in a 6-well plate (2 x 10^5^ cells per well) and allowed to adhere overnight. Cells were either mock-infected or infected with *C. burnetii* in 0.5 ml RPMI 10% FBS for 2 h. Infected cells were washed extensively with PBS and incubated in 1 ml of fresh growth medium. At 24 hpi, cells were stimulated or not with 100 ng/ml mouse IL-17 recombinant protein (R&D Systems), and at 48 hpi, the supernatant was collected and centrifuged at 15,000 × *g* for 10 min at 4°C for debris/bacterial removal. Cytokine/chemokine concentrations were quantitated using the Mouse Cytokine/Chemokine 44-Plex Discovery Assay® Array- MD44 (Eve Technologies).

### Neutrophil migration assay

Permeable 24-well transwell inserts with a pore size of 8 μm (Corning Costar) were used to study the migration of differentiated MPRO clone 2.1 cells and primary mouse neutrophils. WT, *il-17ra* and *Δtraf6* MH-S cells were plated in a 6-well plate (2×10^5^ cells per well), allowed to adhere overnight, and then mock-infected or infected with WT or *ΔdotA* mutant or *cbu2013::Tn C. burnetii* in 0.5 ml growth medium, for 2 h. Infected cells were washed extensively with PBS, and incubated in 1 ml growth medium. At 24 hpi, cells were stimulated or not with 100 ng/ml of mouse IL-17 recombinant protein (R&D Systems). At 48 hpi the supernatant was collected and centrifuged at 15,000 x *g* for 10 min for debris removal. The differentiated MPRO cells (1×10^6^) or primary mouse neutrophils (1×10^5^) were resuspended in 250 μl serum-free media and added in the insert. 650 μl of conditioned media (cell supernatant) was added to the bottom of each well, and then the inserts were carefully lowered into the wells. RPMI supplemented with 10% FBS with or without 100 ng/ml of recombinant mouse CXCL2/MIP-2 protein (R&D Systems) was used as positive and negative controls, respectively. Neutrophils were incubated for 1 h 30 min at 37 °C in 5% CO₂, after which cells that had migrated through the inserts were collected from the lower chamber.

The collected cell suspension was centrifuged (250 × g, 5 min), and the resulting pellet was resuspended in 200 µl of media prior to cell counting using a Trypan Blue exclusion assay. The percentage of migrated cells was calculated as the ratio of counted cells to the number of cells in reference well, multiplied by 100. Percentage of migrated cells was compared to the corresponding untreated controls.

### Pulldown assays

HEK293T cells were seeded at a density of 2×10^5^ cells per well in a 6-well plate and transfected with FLAG-TRAF6, a gift from John Kyriakis (Addgene plasmid # 21624)^53^ and with mStayGold, EmcB-mStayGold and EmcB-mStayGold (C156A) using Fugene6 transfection reagent (Promega). After 48 h, cells were collected, lysed on ice for 30 min in immunoprecipitation (IP) buffer (15 mM Tris pH 7.4, 150 mM NaCl, 1% Triton X-100, 1 x protease inhibitor cocktail (CST), and cell debris pelleted at 15,000xg, 4°C for 10 min. The supernatant was incubated with anti-FLAG agarose magnetic beads (Thermo Fisher Scientific) at 4°C overnight, washed with the IP buffer, and boiled in laemmli sample buffer to elute bound proteins. The proteins analyzed using SDS-PAGE followed by immunoblotting with anti-FLAG (1:1000 Sigma-Aldrich F1804), anti-ubiquitin (1:1000 Cell Signaling 43124). For endogenous immunoprecipitation, alveolar macrophages were infected with *C. burnetii* for 48 hours, lysed using NP-40 buffer (1% NP-40, 50 mM Tris-HCl, pH7.4, 50 mM EDTA, 150 mM NaCl), and clarified lysates incubated with anti-TRAF6 antibody (Santa Cruz Biotechnology sc-8409) and protein G beads (Thermo Fisher 88847) overnight at 4 °C. Beads were washed with the IP buffer, beads were then boiled in laemmli sample buffer to elute bound proteins. The proteins were analyzed using SDS-PAGE followed by immunoblotting with anti-TRAF6 (1:1000 Abcam Anitbodies ab33915), anti-ubiquitin (1:1000 Cell Signaling 43124).

### Plasmids

For mammalian ectopic expression, wildtype of C156A *cbu2013* (NC_002971.4:c1921006-1919909) was cloned into the pTwist CMV BG WPRE Neo vector backbone, (Twist Biosciences). The *C. burnetii* complementation plasmid was generated by cloning *cbu201* into pJB-2HA-CBU2013 using forward primer 5′-AGATTACGCTGTCGACATGCCATCGATAAATTTGACGACG-3′ and reverse primer 5′-GCATGCCTCAGTCGATTAAGAAACTAGCTGAAGATGAGGATTTGAGT-3′.

The resulting PCR products were ligated into the SalI-linearized pJB-Kan-p1169-2xHA plasmid using In-Fusion HD cloning (Takara Bio). Plasmid constructs were validated through Sanger didexoy sequencing (ACGT) and Oxford Nanopore Technology (Plasmidsaurus)

## Data analyses

Image processing and analyses were done in ImageJ (Fiji) software. Statistical analyses were performed as indicated in the figure legends using Prism (GraphPad).

## Acknowledgments

We thank Seth Winfree, Rey Carabeo, and members of the Gilk Lab for helpful discussions and critical feedback. We also thank Matteo Bonazzi for providing the *cbu2013::Tn* mutant. This research was supported by National Institutes of Health (1R21AI149723 to SDG), NIH/NIGMS (T32GM153375 to AS) and an American Heart Association Postdoctoral Fellowship (834525 to TMC). We have no conflicts of interest to declare.

## Figure legends

**Supplementary Figure 1. Depletion of il-17ra, traf6 and ACT1 in MH-S cells**. **(A-B)** Quantitative expression of il-17ra and traf6 in MH-S cells by qRT-PCR using specific primers confirmed knockout of il-17ra and traf6 using CRISPR-Cas9. **(C)** Immunoblot showing depletion of ACT1 in MH-S using siRNA. Control cells were transfected with non-targeting siRNA (NT).

**Supplementary Figure 2. IL-17 increases NOX activity and ROS production in alveolar macrophages. (A)** WT and Δ*il-17ra* MH-S cells were treated or not with recombinant IL-17 (100 ng/ml) and the NOX activity was measured. IL-17 stimulation significantly increased NOX activity WT cells, but not in *il-17ra* cells. Data shown as means ± SEM from three independent experiments. Statistical significance was determined by one-way ANOVA with Tukey’s multiple comparisons test, ***p<0.001. **(B)** ROS levels in Δ*il-17ra* were measured using CellROX Green. Data are shown as the mean ± SEM of at least 20 cells per condition in each of three independent experiments. Statistical significance was determined by two-tailed unpaired student’s t-test.

**Supplementary Figure 3. IL-17 treatment does not affect the secretion of non-specific cytokines. (A)** Migration Assay of MPRO cells incubated with the supernatant of mock- and WT-infected Δ*il-17ra* cells treated with IL-17 did not show any difference in neutrophil migration. **(B)** Primary neutrophils isolated from C57BL/6 bone marrow cells for transwell migration assay and the purity validated using flow cytometry (∼75% purity).

