## Supplementary figures and images for "*Coxiella burnetii* deubiquitinates host TRAF6 to modulate the macrophage innate immune response"

### Supplemental Figures

Supplementary Figure 1

A

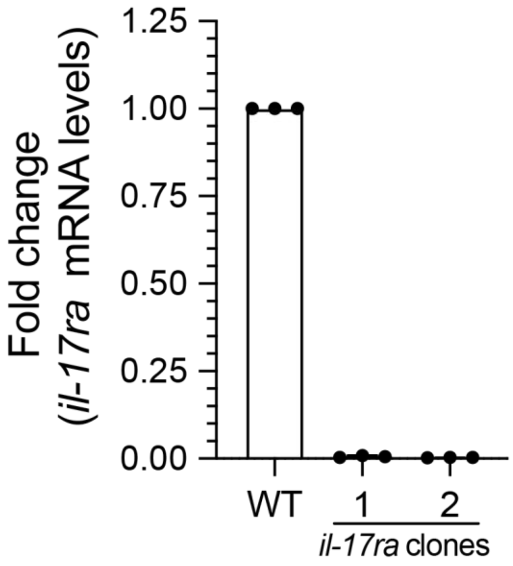


B

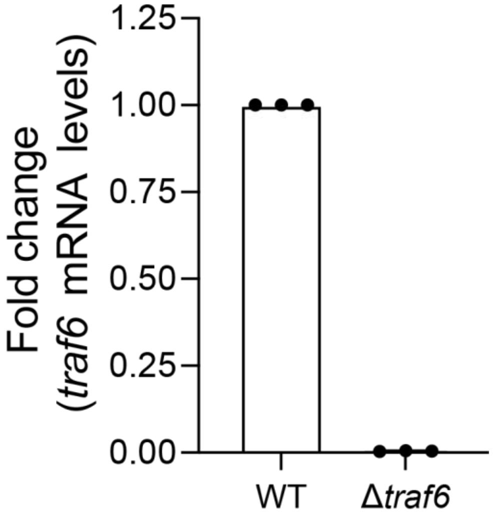


C

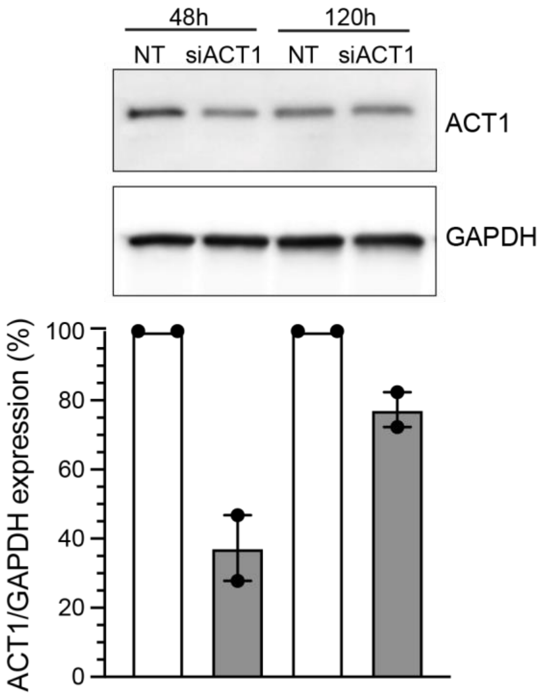

Supplementary Figure 2

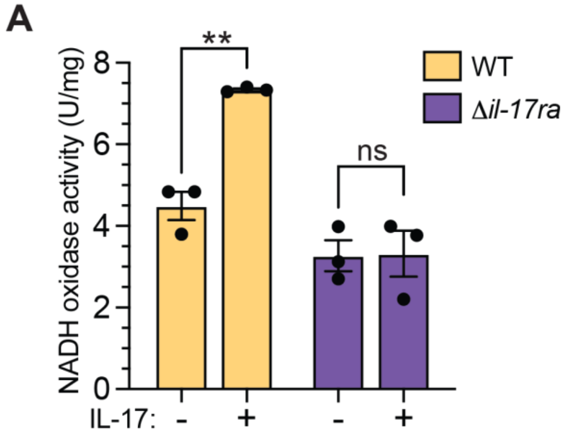


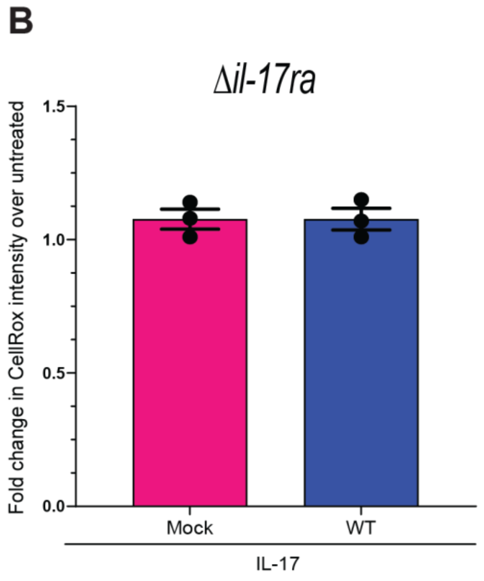


Supplementary Figure 3

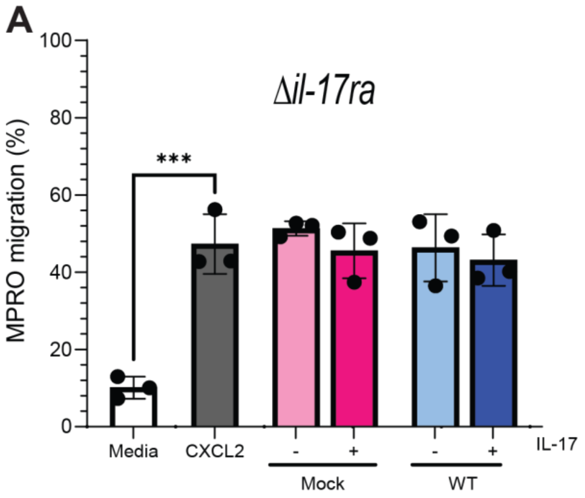


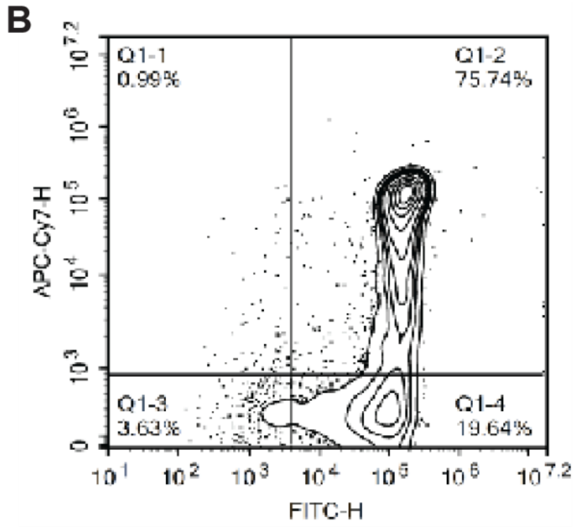
